# The Biogeography of Coelurosaurian Theropods and its Impact on their Evolutionary History

**DOI:** 10.1101/634170

**Authors:** Anyang Ding, Michael Pittman, Paul Upchurch, Jingmai O’Connor, Daniel J. Field, Xing Xu

## Abstract

The Coelurosauria are a group of mostly feathered theropods that gave rise to birds, the only dinosaurs that survived the Cretaceous-Paleogene extinction event and are still found today. Between their first appearance in the Middle Jurassic up to the end Cretaceous, coelurosaurs were party to dramatic geographic changes on the Earth’s surface, including the breakup of the supercontinent Pangaea, and the formation of the Atlantic Ocean. These plate tectonic events are thought to have caused vicariance or dispersal of coelurosaurian faunas, influencing their evolution. Unfortunately, few coelurosaurian biogeographic hypotheses are supported by quantitative evidence. Here, we report the first, broadly-sampled quantitative analysis of coelurosaurian biogeography using the likelihood-based package BioGeoBEARS. Mesozoic geographic configurations and changes are reconstructed and employed as constraints in this analysis, including their associated uncertainties. We use a comprehensive time-calibrated coelurosaurian evolutionary tree produced from the Theropod Working Group phylogenetic data matrix. Six biogeographic models in the BioGeoBEARS package with different assumptions about the evolution of spatial distribution are tested against the geographic constraints. Our results statistically favour the DIVALIKE+J and DEC+J models, which allow vicariance and founder events, supporting continental vicariance as an important factor in coelurosaurian evolution. Ancestral range estimation indicates frequent dispersal events via the Apulian Route (connecting Europe and Africa during the Early Cretaceous) and the Bering Land Bridge (connecting North America and Asia during the Late Cretaceous). These quantitative results are consistent with commonly inferred Mesozoic dinosaurian dispersals and continental-fragmentation-induced vicariance events. In addition, we recognise the importance of Europe as a dispersal centre and gateway in the Early Cretaceous, as well as other vicariance events like those triggered by the disappearance of land-bridges.

## INTRODUCTION

Coelurosauria is a derived clade of theropod dinosaurs which includes Tyrannosauroidea, Compsognathidae, Ornithomimosauria, Alvarezsauroidea, Therizinosauroidea, Oviraptorosauria, Dromaeosauridae, Troodontidae and Avialae (Brusatte, et al., 2014). A large portion of coelurosaurs were feathered, while some of them, mainly from the Avialae, acquired powered flight ability (Xu, et al., 2014). Most coelurosaurian lineages lived from the Middle Jurassic to the end of the Cretaceous, with only a subset of avian taxa (Neornithes) surviving the Cretaceous-Paleogene (K-Pg) extinction event (Xu, et al., 2014). We use Avialae here following the preferred term used by most non-avian theropod workers, whereas Aves is preferred by most ornithologists. During the late Mesozoic, coelurosaurs and other dinosaurs lived through dramatic geographic changes (Upchurch, et al., 2002a): plate tectonic activity caused continents to break apart to form new oceans and seas, produced intermittent re-connections, and prompted fluctuations in sea level which further modified palaeogeographic relationships (Condie, 2013). These geographic configurations and changes are presumed to have affected coelurosaur populations and faunas, impacting their pattern and tempo of evolution (Benton and Harper, 2013). Evaluating this impact is crucial if we are to fully understand the most significant events in coelurosaurian evolution, including the acquisitions of herbivory and early theropod flight.

Biogeographic studies focus on the geography-dependent processes that lead to alterations in faunal distributions and speciation (Benton and Harper, 2013). Four of the most fundamental biogeographic processes are (Sereno, 1999a; Upchurch, et al., 2002a; Isabel, et al., 2004; Sanmartín and Ronquist, 2004): 1. *dispersal*: when a fauna expands its distribution range, 2. *regional extinction*: when the distribution shrinks, 3. *sympatry*: when speciation happens within the ancestral distribution range of the fauna, and 4. *vicariance*: when speciation takes place due to the separation of two populations by a geographic barrier. These processes play important roles in organismal evolution and are sensitive to geographic conditions.

Numerous biogeographic hypotheses have been proposed for clades within Coelurosauria, though the vast majority of these are narratives (by the definition of Ball (1975:) because they tend to read the fossil record literally. A few studies (e.g. Loewen, et al. (2013:) have applied quantitative phylogenetic biogeographic analyses to groups such as Tyrannosauroidea, but the majority of coelurosaurian subclades and the group as a whole have not been investigated using such approaches. At present, therefore, much of our knowledge of coelurosaurian biogeographic history comes from studies of Dinosauria as a whole (e.g. Upchurch, et al. (2002b; O’Donovan, et al. (2018:). To address these deficits, we perform the first quantitative biogeographic analysis focused on the Coelurosauria as a whole.

## Mesozoic Palaeogeography

The palaeogeography of Pangaea provides an important backdrop to the evolution of coelurosaurs, and information on this topic is required in order to support the geographic constraints we apply in our biogeographic analyses. Below, therefore, we briefly outline key aspects of Mesozoic palaeogeography.

The Mesozoic witnessed the breakup of the supercontinent Pangaea and the establishment of global geography close to the modern arrangement (Scotese, 2001). However, narrow land bridges connecting isolated landmasses did appear during short time intervals, and shallow epicontinental seas existed throughout the Mesozoic, especially within Laurasian landmasses (Poropat, et al., 2016).

During the early Mesozoic all continents were joined together to form Pangaea, although the Laurasia-Gondwana connection was only present between North America and (Africa + South America) (Smith, et al., 2004). The breakup of Pangaea initiated during the Middle Jurassic, starting with the separation of North America from South America, together with the opening of the northern Atlantic (Bardet, et al., 2014). The complete separation of Laurasia and Gondwana dates back to the Kimmeridgian stage of the Late Jurassic (Gaina, et al., 2013). Rifting and sea floor spreading between Africa, Indo-Madagascar, and Antarctica began later, during the Tithonian (Seton, et al., 2012). The Turgai Sea existed between Asia and Europe throughout the late Mesozoic (especially the Late Cretaceous), although intermittent land connections occurred because of sea level fluctuations (Baraboshkin, et al., 2003; Smith, et al., 2004). During the Late Jurassic and earliest Cretaceous, Gondwana gradually separated into two large continents comprising South America + Africa (Samafrica) and Antarctica + Indo-Madagascar + Australia (East Gondwana) (Eagles and König, 2008). However, the sequence and timing of the breakup of Gondwana remains controversial (e.g. Sereno, et al. (2004; Krause, et al. (2006; Krause, et al. (2007; Upchurch (2008; Ali and Krause (2011:) and several workers have proposed that South America and Antarctica maintained a contact via Patagonia and the West Antarctic Peninsula throughout some or all of the Cretaceous (see review in Poropat, et al. (2016:). During the earliest Cretaceous, the Apulian Route was established (Zarcone, et al., 2010). This connection between south-western Europe and north-western Africa was the first between Laurasia and Gondwana after the breakup of Pangaea (Ezcurra and Agnolín, 2012). The land connection between eastern North America and Western Europe finally disappeared with the full establishment of the north Atlantic Ocean in the Barremian or Aptian (Seton, et al., 2012). Later in the late Aptian and Albian stages, the Bering Land Bridge connected north-eastern Asia and north-western North America for the first time (Plafker and Berg, 1994). This land bridge was probably absent during the Cenomanian-Santonian, but was potentially re-established in the late Campanian and perhaps the Maastrichtian (Brikiatis, 2014). The Western Interior Seaway separating North America into eastern and western portions (known as Appalachia and Laramidia respectively) was present from then on throughout the Late Cretaceous until a possible reconnection during the Maastrichtian (Smith, et al., 2004; Farke and Phillips, 2017). Africa and South America separated from each other at the end of the Albian Stage, after the isolation of Indo-Madagascar during the Aptian Stage (Eagles and König, 2008). India separated from Madagascar during the latest Cretaceous (Plafker and Berg, 1994). By the end of the Cretaceous, global geography had a configuration that resembles the modern one, though Africa and India did not collide with Eurasia, and the Patagonia-Antarctica connection might not have been severed until the Cenozoic (Matthews, et al., 2016).

## Geographic and Temporal Distribution of Coelurosaurs

Most known fossil coelurosaurs are from Laurasia (1083 occurrences recorded at the time of writing in the Paleobiology Database; https://paleobiodb.org/), with only a few occurrences in Gondwana (59 recorded in the Paleobiology Database). Currently, the earliest known coelurosaurs are the proceratosaurids *Proceratosaurus* (von Huene, 1926) and *Kileskus* (Averianov, et al., 2010) from the Bathonian stage of the Middle Jurassic of southern England and central Russia respectively. The most stemward coelurosaurs are *Bicentenaria* from Argentina (Novas, et al., 2012), *Coelurus* from America (Marsh, 1879), and *Zuolong* from China (Choiniere, et al., 2010a). The occurrences of tyrannosauroids during the Middle Jurassic and basal paravians during the early Late Jurassic (e.g. *Anchiornis*) imply that major lineages of coelurosaurs were established by the Middle-Late Jurassic (Rauhut, et al., 2010; Choiniere, et al., 2012). Some authors have argued that the clades, including Compsognathidae, Tyrannosauroidea, and Maniraptoriformes, probably originated during or even before the Middle Jurassic (Rauhut, et al., 2010), so predating the separation of Laurasia and Gondwana. The currently known geographic and temporal distributions of the major coelurosaurian clades Tyrannosauroidea, Compsognathidae, Ornithomimosauria, Alvarezsauroidea, Therizinosauroidea, Oviraptorosauria, Dromaeosauridae, Troodontidae, and Avialae are discussed in more detail below:

### Tyrannosauroidea

Tyrannosauroids include the infamous *Tyrannosaurus rex* and its closest relatives (Brusatte, et al., 2010). Stemward Jurassic tyrannosauroids had a wide distribution in Laurasia as indicated by *Guanlong* from Asia (Xu, et al., 2006), *Juratyrant* from Europe (Brusatte and Benson, 2013), and *Stokesosaurus* from North America (Benson, 2008). Dispersal events within Laurasian landmasses are inferred during that period of time (Rauhut, et al., 2010). More crownward tyrannosaurids are mostly known from the Late Cretaceous of Asia and western North America (Brusatte, et al., 2010). The existence of closely-related taxa in both Asia and western North America just before the end of the Cretaceous, as in other coelurosaur clades, may suggest faunal exchange events between these landmasses at that time (Brusatte, et al., 2010). Traditionally, it was thought that tyrannosaurs were restricted to Laurasian landmasses, including North America, Europe, and Asia, but Gondwanan material challenges this (Benson, et al., 2010). The Australian occurrence of a possible tyrannosauroid is inferred based on a late Early Cretaceous pubis described in 2010 (Benson, et al.). Some authors (Porfiri, et al., 2014; Porfiri, et al., 2018) put megaraptorids within Tyrannosauroidea, which may infer a wider distribution of the clade within Gondwanan landmasses.

### Compsognathidae

Compsognathids are comparatively small stemward coelurosaurs known from the Late Jurassic to Early Cretaceous (Hwang, et al., 2004). Laurasian compsognathids occur in North America (Osborn, 1903), Europe (Göhlich and Chiappe, 2006), and Asia (Hwang, et al., 2004) but only one Gondwanan taxon, *Mirischia*, from the Albian of South America is known (Naish, et al., 2004). Since *Mirischia* is the youngest and most crownward compsognathid, a dispersal event from Laurasia to Gondwana seems likely, most probably from Europe to South America, via Africa (Naish, et al., 2004). If this occurred, more crownward compsognathids are expected to be found in Africa in the future.

### Ornithomimosauria

The slender ‘ostrich-like’ ornithomimosaurians lived during the Cretaceous period (Xu, et al., 2011a). Ornithomimosaurians are found in all Laurasian landmasses, with most frequent occurrences in Asia (Xu, et al., 2011a). *Nqwebasaurus* from South Africa is the most stemward ornithomimosaurians and is also the only one from Gondwana (De Klerk, et al., 2000). Given the close relationship between *Nqwebasaurus* and stemward Laurasian ornithomimosaurians, the clade is inferred to have achieved a wide distribution before the breakup of Pangaea (Allain, et al., 2014). The crownward North American ornithomimosaurians, including *Ornithomimus* and *Struthiomimus*, form a monophyletic group (Xu, et al., 2011a). This phylogenetic and geographic pattern has been explained by a single dispersal event from Asia to North America via the Bering Land Bridge during the latest Cretaceous (Ji, et al., 2003; Jin, et al., 2012).

### Alvarezsauroidea

Alvarezsauroids are known for their small, crownward forms that have especially large first fingers (Xu, et al., 2011b). Until recently, alvarezsauroids were only known from the Late Cretaceous, including intermediate forms from South America (e.g. *Alvarezsaurus* (Bonaparte, 1991) and more crownward forms from Asia and North America (e.g. *Mononykus* (Altangerel, et al., 1993) and *Albertonykus* (Longrich and Currie, 2009b)). This led to a South American origin being proposed for the clade (Longrich and Currie, 2009b). The discovery of the stem alvarezsauroid *Haplocheirus* from the Late Jurassic of China overturned this origin hypothesis (Choiniere, et al., 2010b). On the basis of a quantitative analysis (Xu, et al., 2011b), it was later proposed that if alvarezsauroids originated in central Asia shortly before the breakup of Pangaea, a dispersal from Asia to South America probably occurred before the Late Cretaceous, most likely via Europe and Africa. A distal tibiotarsus from Romania is the only suspected record of the group in Europe (Naish and Dyke, 2004), but no African records are yet known. The dispersal of Patagonian alvarezsaurids to Asia has become a consensus recently (Xu, et al., 2011b; Averianov and Sues, 2017). An additional Late Cretaceous dispersal event from Asia to North America is inferred to explain the occurrence of the crownward North American alvarezsaurid *Albertonykus* which is closely related to crownward Asian forms (Longrich and Currie, 2009b; Agnolin, et al., 2012).

### Therizinosauria

Therizinosaurians are a Cretaceous coelurosaur clade that evolved herbivory, as also seen in Ornithomimosauria and Oviraptorosauria (Zanno and Makovicky, 2011). Most therizinosaurians are from the Cretaceous of Asia, especially China and Mongolia (Zanno, 2010). The most stemward therizinosaurian, *Falcarius* from the Barremian of Utah, potentially indicates a vicariance event resulting from the separation of North America and Asia during the Early Cretaceous, or a dispersal of stemward therizinosaurians from North America to Asia via the controversial land connections proposed across the proto-Atlantic and Turgai Sea (Zanno, 2010). More fossil evidence, such as earlier and/or confirmed European records are required to address this issue further. The other non-Asian therizinosaurians are crownward forms from the early Late Cretaceous of North America (e.g. *Nothronychus*) whose ancestors potentially dispersed from Asia via the Bering Land Bridge during its establishment in the later stages of the Early Cretaceous (Kirkland and Wolfe, 2001; Zanno, 2010; Fiorillo and Adams, 2012). This dispersal event receives further support in the form of a potential therizinosaurian track found in Alaska, USA, which is one side of the modern Bering Strait (Fiorillo and Adams, 2012; Fiorillo, et al., 2018).

### Oviraptorosauria

Oviraptorosaurians are known for the preservation of evidence of their brooding behaviour, and include crownward forms with short, elaborate skulls (Clark, et al., 2001). Stemward oviraptorosaurians, including *Incisivosaurus*, caudipterygids and *Avimimus*, are solely Asian taxa that lived before the Late Cretaceous (Funston and Currie, 2016). More crownward taxa have parrot-like beaks, with or without bony skull crests and have been divided into two subclades, Caenagnathidae and Oviraptoridae (Lü, et al., 2015). While known oviraptorids are restricted to Asia, the Caenagnathidae include both North American and Asian taxa (Xu, et al., 2007; Funston and Currie, 2016). The presence of the Albian-aged *Microvenator* in North America is probably attributable to a dispersal event of stemward oviraptorosaurs from Asia via the Bering Land Bridge (Makovicky and Sues, 1998). Like several other coelurosaurian clades, the Late Cretaceous caenagnathids spread across Asia and North America (Funston and Currie, 2016).

### Dromaeosauridae

Dromaeosaurids together with troodontids are the closest relatives of birds (Turner, et al., 2012). Dromaeosaurids and troodontids have a hyper-extendable second toe, while dromaeosaurids include taxa with rod-like tails comprising caudal vertebrae bound by elongated prezygapophyses (Turner, et al., 2012). Dromaeosaurids have a broad geographic distribution across Laurasia and Gondwana throughout the Cretaceous (Turner, et al., 2012). Laurasian taxa include stemward forms such as *Mahakala* (Turner, et al., 2007) as well as crownward ones like the renowned *Velociraptor* (Osborn, et al., 1924). Stemward dromaeosaurids from Gondwanan landmasses, including *Rahonavis, Buitreraptor, Neuquenraptor* and *Austroraptor* form a single clade (Turner, et al., 2012). This may indicate a vicariance event due to the separation of Laurasia and Gondwana during the Middle Jurassic (Makovicky, et al., 2005; Novas and Pol, 2005). Antarctic occurrences of dromaeosaurids were also inferred by some pedal fossil fragments, which, together with other Gondwanan taxa, might imply a cosmopolitan distribution of the clade before the breakup of Pangaea (Case, et al., 2007). Although Jurassic teeth from Laurasia have been referred to Dromaeosauridae (Goodwin, et al., 1999; Vullo, et al., 2014), more substantial fossil evidence is needed to confirm this important early record. The establishment of the Bering Land Bridge during the later stages of the Early Cretaceous and again in the latest Cretaceous, has been proposed as a potential explanation of the flourishing of Velociraptorinae in Asia and the occurrence of microraptorines (*Hesperonychus*) in North America (Longrich and Currie, 2009a; Turner, et al., 2012). Faunal exchange between Europe and Asia has also been inferred during the Cretaceous based on the close relationship between the European dromaeosaurid *Balaur* and other Laurasian dromaeosaurids (Csiki, et al., 2010; Brusatte, et al., 2013). Whilst flight capabilities have been proposed in the microraptorine *Microraptor*, these relate to relatively short distance flight and probably did not affect the dispersal ability of dromaeosaurids over continental scales, but perhaps came into play in archipelago settings (Chatterjee and Templin, 2007).

### Troodontidae

Troodontids can be distinguished from dromaeosaurids by their numerous, closely-packed teeth (Currie, 1987). Most of these close avian relatives are from Asia (Lü, et al., 2010). North American occurrences of the clade are restricted to *Geminiraptor* from the Early Cretaceous of Utah, USA (Senter, et al., 2010) and several crownward taxa from the Late Cretaceous (Leidy, 1856; Zanno, et al., 2011). While the occurrence of crownward North American troodontids, represented by *Troodon*, can be attributed to a dispersal from Asia via the Bering Land Bridge during the Campanian and Maastrichtian stages of the Late Cretaceous (Dodson, et al., 2004), *Geminiraptor* and abundant teeth referred to troodontids from Europe indicate that multiple dispersal events might have happened within Laurasia even before the Late Cretaceous (Senter, et al., 2010). The first Gondwanan troodontid was reported based on a tooth found in the Late Cretaceous of India, with this occurrence reflecting either a dispersal event from Laurasia, or a much wider distribution of the clade before the breakup of Pangea (Goswami, et al., 2013).

### Avialae

This clade includes early birds and their modern descendants (Padian, 2004) and by the Late Cretaceous avialans had achieved a global geographic distribution (Brocklehurst, et al., 2012). The controversial Late Jurassic stem paravians, the anchiornithids, proposed as both stem birds and troodontids were previously only known from Liaoning, China (Godefroit, et al., 2013), but have recently been confirmed in Europe (*Ostromia*) (Foth and Rauhut, 2017). This might indicate a Late Jurassic dispersal event from Asia to Europe (Foth and Rauhut, 2017) given that *Archaeopteryx*, the most widely accepted oldest and most stemward bird, is from Germany (Wellnhofer, 2009; Foth, et al., 2014). The second oldest avifaunas are in the Early Cretaceous Hauterverian - Barremian, in which the more crownward clades Ornithuromorpha and Enantiornithes first appear, are found in China (Zhou and Zhang, 2006) and Mongolia (O’Connor and Zelenkov, 2013; Zelenkov and Averianov, 2016) where both clades are represented, and in Spain, where only enantiornithines have been found (Sanz, 1990; Sanz, et al., 1995; Sanz, et al., 1996). Only the Hauterverian – Aptian Jehol Biota preserves non-ornithothoracines (*Jeholornis* with its long bony tail and stemward pygostylians *Sapeornis* and *Confuciusornis*) (Zhou and Zhang, 2006). Slightly younger deposits find enantiornithines in an even wider distribution, present in Gondwanan deposits in Brazil (de Souza Carvalho, et al., 2015) and Australia (Close, et al., 2009), with the earliest hesperornithiforms preserved in late Albian deposits in the UK (Galton and Martin, 2002). During the Late Cretaceous, enantiornithines and crownward ornithuromorphs (ornithurines) have a global distribution, with records in Asia (e.g. *Gobipteryx*), South America (e.g. *Patagopteryx*), North America (e.g. *Ichthyornis*), Europe (e.g. *Baptornis*), Madagascar (*Vorona*) (Elzanowski, 1974; Martin and Bonner, 1977; Alvarenga and Bonaparte, 1992; Forster, et al., 1996; Clarke, 2004) and Antarctica (*Vegavis*) (Clarke, et al., 2005).

Theropod flight appeared in the Middle or Late Jurassic at the latest (Xu, et al., 2014), but the exact time or times when powered flight was acquired is still under debate (Brocklehurst, et al., 2012; Allen, et al., 2013; Zheng, et al., 2013; Xu, et al., 2014; Dececchi, et al., 2016). Modern avian flight ability varies widely from species to species (Tobalske, et al., 2003) with some birds being flightless whilst others like terns (*Sterna*) are capable of migrating across oceans (Tobalske, et al., 2003). Given the general absence of functional-informative soft tissue evidence in avian fossils, the dispersal ability of Mesozoic avians is even harder to estimate than modern birds, providing major challenges to biogeographic analysis of this clade. However, the distribution of Late Cretaceous taxa such as the enantiornithine *Martinavis*, found in North and South America and in Europe (Walker, et al., 2007), may suggest at least some taxa were able to migrate long distances, and were unrestricted in their dispersal relative to non-avian dinosaurs.

## Major Coelurosaurian Biogeographic Hypotheses

Besides the clade-level biogeographic hypotheses summarised above (see *Geographic and temporal distribution of coelurosaurs* section), analyses of dinosaur biogeography as a whole, including coelurosaurs, have given different emphasis to particular biogeographic processes.

Many authors attach particular importance to vicariance events because of the global continental fragmentation that occurred during the late Mesozoic (Sereno, 1999b; Upchurch, et al., 2002b; Choiniere, et al., 2012). The Middle Jurassic occurrences of tyrannosauroids (e.g. *Kileskus* and *Proceratosaurus*) and Late Jurassic avian occurrences (e.g. *Aurornis* and *Anchiornis* [but also proposed as troodontids]), are consistent with the idea that major coelurosaurian lineages were established at least before the Late Jurassic (Rauhut, et al., 2010; Choiniere, et al., 2012). Together with several Gondwanan stemward coelurosaurian occurrences (e.g. *Bicentenaria* from South America and *Nqwebasaurus* from Africa), a geographically widespread distribution of coelurosaurian lineages before the breakup of Pangaea has been inferred, which makes vicariance possible upon separation of the continents (Choiniere, et al., 2012). Proposed continental scale vicariance events include the Laurasia-Gondwana separation during the Late Jurassic (as shown in Fig. 1; Hypothesis 1) and the final disconnection of South America and Africa during the Early Cretaceous (as shown in Fig. 2; Hypothesis 2) (Sereno, 1999b). Possible vicariance-induced phylogenetic patterns have been identified in the distributions of maniraptoran lineages (Makovicky, et al., 2005) and Ornithomimosauria (De Klerk, et al., 2000). However, others see continental-scale vicariance as a rare occurrence, and argue that regional extinctions were primarily responsible for late Mesozoic dinosaurian distributions, based on the observation that vicariance-like repeated area relationships can also be explained by regional extinction events and that many clades were widespread early in their evolutionary history and seem to become more geographically restricted subsequently (Sereno, 1997; 1999a; Barrett, et al., 2011; Benson, et al., 2012; Carrano, et al., 2012).

**Fig. 1.**
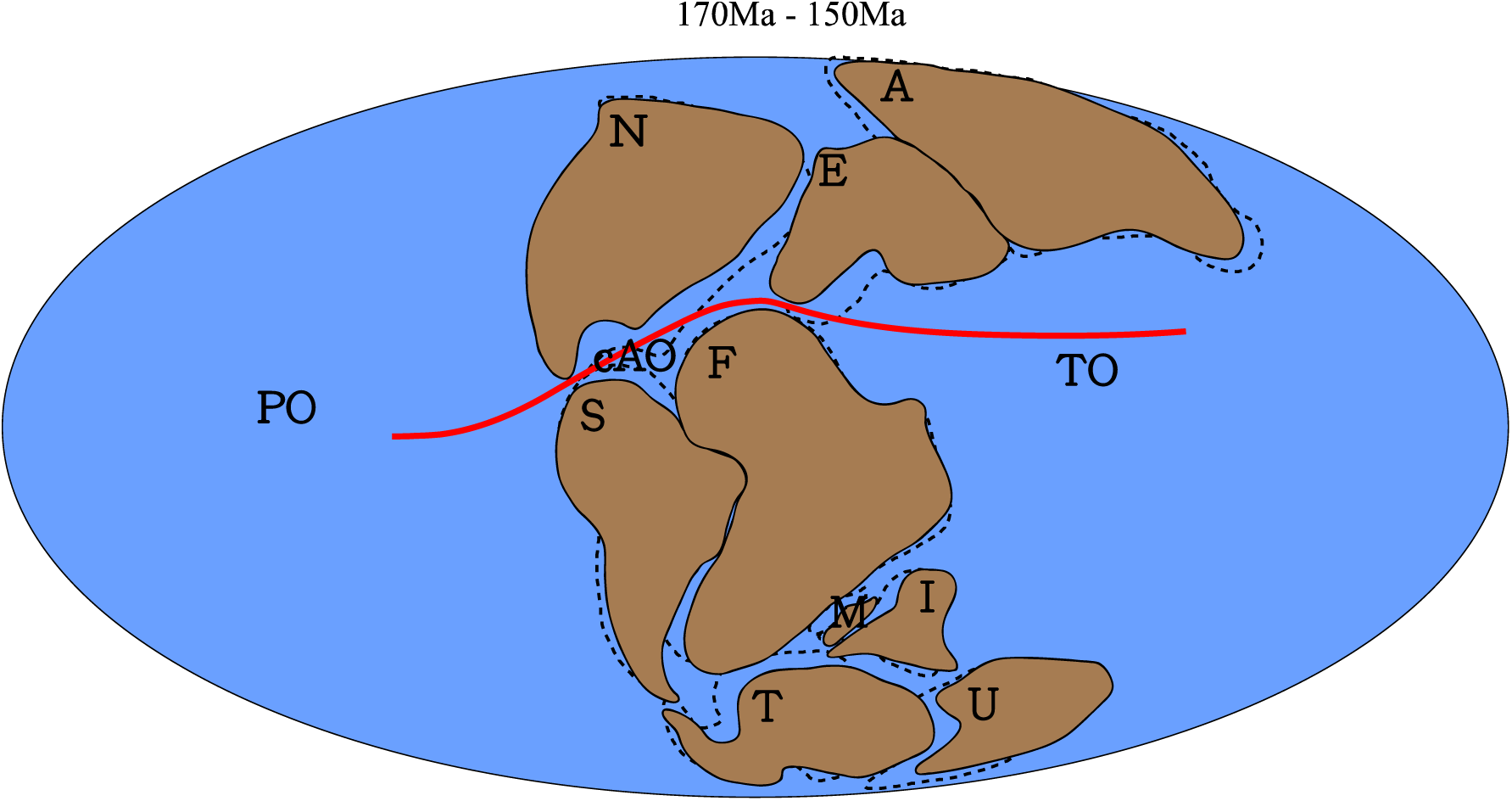
Hypothesis 1, Late Cretaceous Laurasia-Gondwana vicariance event during the Late Jurassic. The separation between Laurasia and Gondwana was established in the Kimmeridgian Stage. The red line denotes the approximate position of the hypothesized biogeographical barrier: the central Atlantic Ocean (cAO) and Tethys Ocean (TO); Dotted lines denote palaeogeography at 170 Ma, whilst solid lines denote it at 150 Ma. Palaeomap after (Matthews, et al., 2016). Abbreviations: A, Asia; cAO, central Atlantic Ocean; E, Europe; F, Africa; I, India; M, Madagascar; N, North America; PO, Pacific Ocean; S, South America; T, Antarctica; TO, Tethys Ocean; U, Australia.

**Fig. 2:**
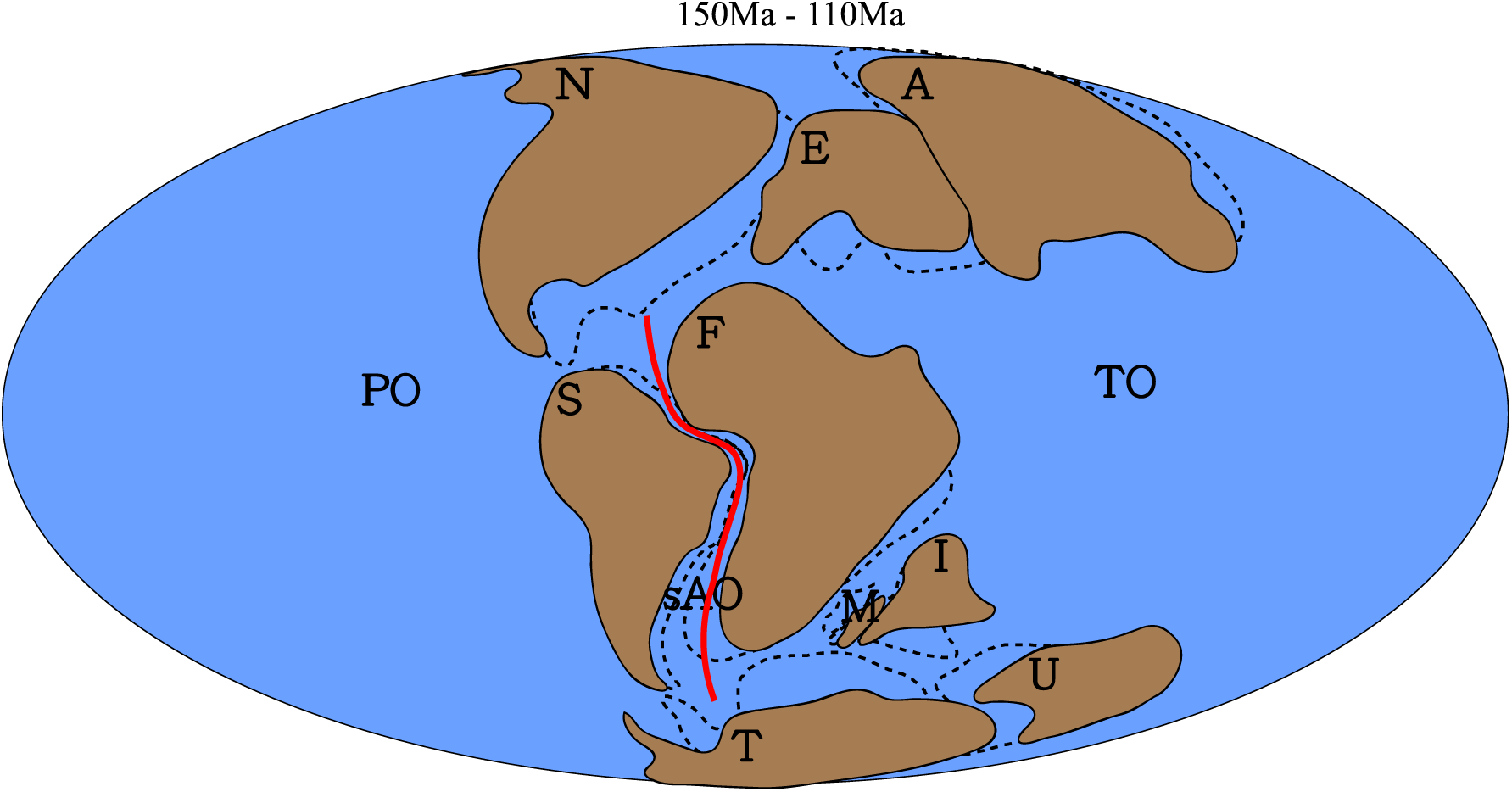
Hypothesis 2, South America-Africa vicariance event during the late Early Cretaceous. The separation between South America and Africa was established in the Albian Stage. The red line denotes the approximate position of the hypothesized biogeographical barrier: the south Atlantic Ocean (sAO); Dotted lines denote paleogeography 150 Ma, whilst solid lines denote it 110 Ma. Palaeomap after (Matthews, et al., 2016). Abbreviations: A, Asia; F, Africa; I, India; M, Madagascar; N, North America; PO, Pacific Ocean; S, South America; sAO, south Atlantic Ocean; E, Europe; T, Antarctic; TO, Tethys Ocean; U, Australia.

Most authors agree that intercontinental dispersal played a key role in creating dinosaurian (coelurosaurian) biogeographic patterns (Sereno, 1999b; Brusatte, et al., 2013; Dunhill, et al., 2016). Such dispersal events are implied from the fossil record and phylogenetic relationships. Faunal dispersal events in both directions via the Early Cretaceous Apulian Route were inferred by different authors: faunal assemblages from the Santana Formation of northern South America that are similar to Laurasian ones suggest possible Asian dinosaur dispersal to Africa via Europe (Naish, et al., 2004); and the presence of Gondwanan faunas in Europe indicates dispersal events in the opposite direction (Ezcurra and Agnolín, 2012; Dal Sasso, et al., 2016). They are unified here as an Africa-Europe faunal exchange hypothesis (Hypothesis 3), as shown in Fig. 3. Frequent dispersal events within coelurosaurian lineages (including Tyrannosauroidea, Therizinosauroidea, and Dromaeosauridae), enabled by the Bering Land Bridge are well documented and accepted (Makovicky and Sues, 1998; Ji, et al., 2003; Dodson, et al., 2004; Longrich and Currie, 2009b; Brusatte, et al., 2010; Zanno, 2010; Turner, et al., 2012), as discussed in the last section (see *Geographic and temporal distribution of coelurosaurs* section) (Fig. 4; Hypothesis 4): this includes impacts from both the Early Cretaceous and Late Cretaceous establishments of the land-bridge.

**Fig. 3.**
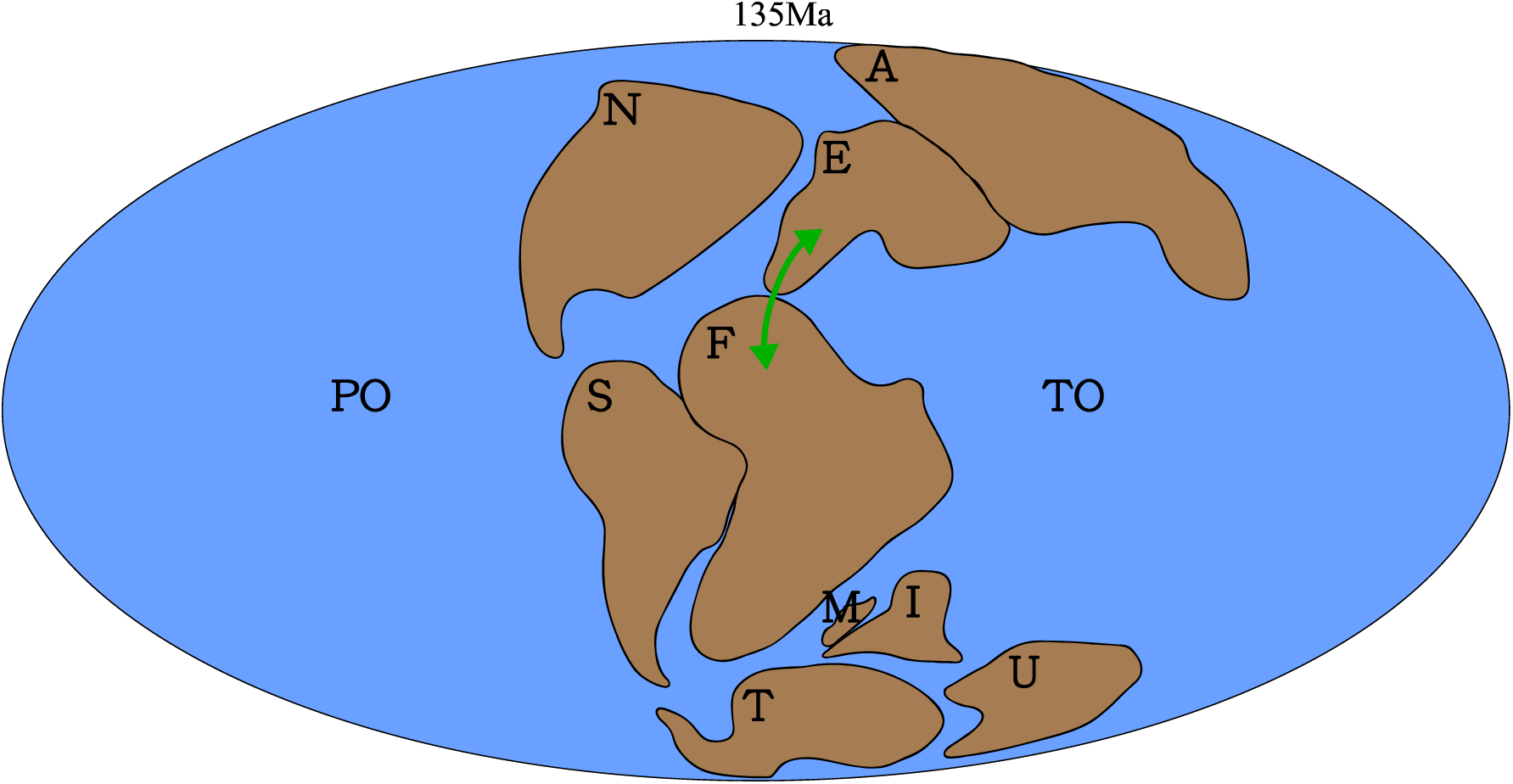
Hypothesis 3, late Early Cretaceous Europe-Africa faunal exchange via the Apulian Route. (Hypothesis 3). The green arrowed lines denote the approximate dispersal directions and dispersal routes; Dotted lines denote palaeogeography Solid lines denote palaeogeography at 135 Ma. Palaeomap after (Matthews, et al., 2016). Abbreviations: A, Asia; E, Europe; F, Africa; I, India; M, Madagascar; N, North America; PO, Pacific Ocean; S, South America; T, Antarctica; TO, Tethys Ocean; U, Australia.

**Fig. 4.**
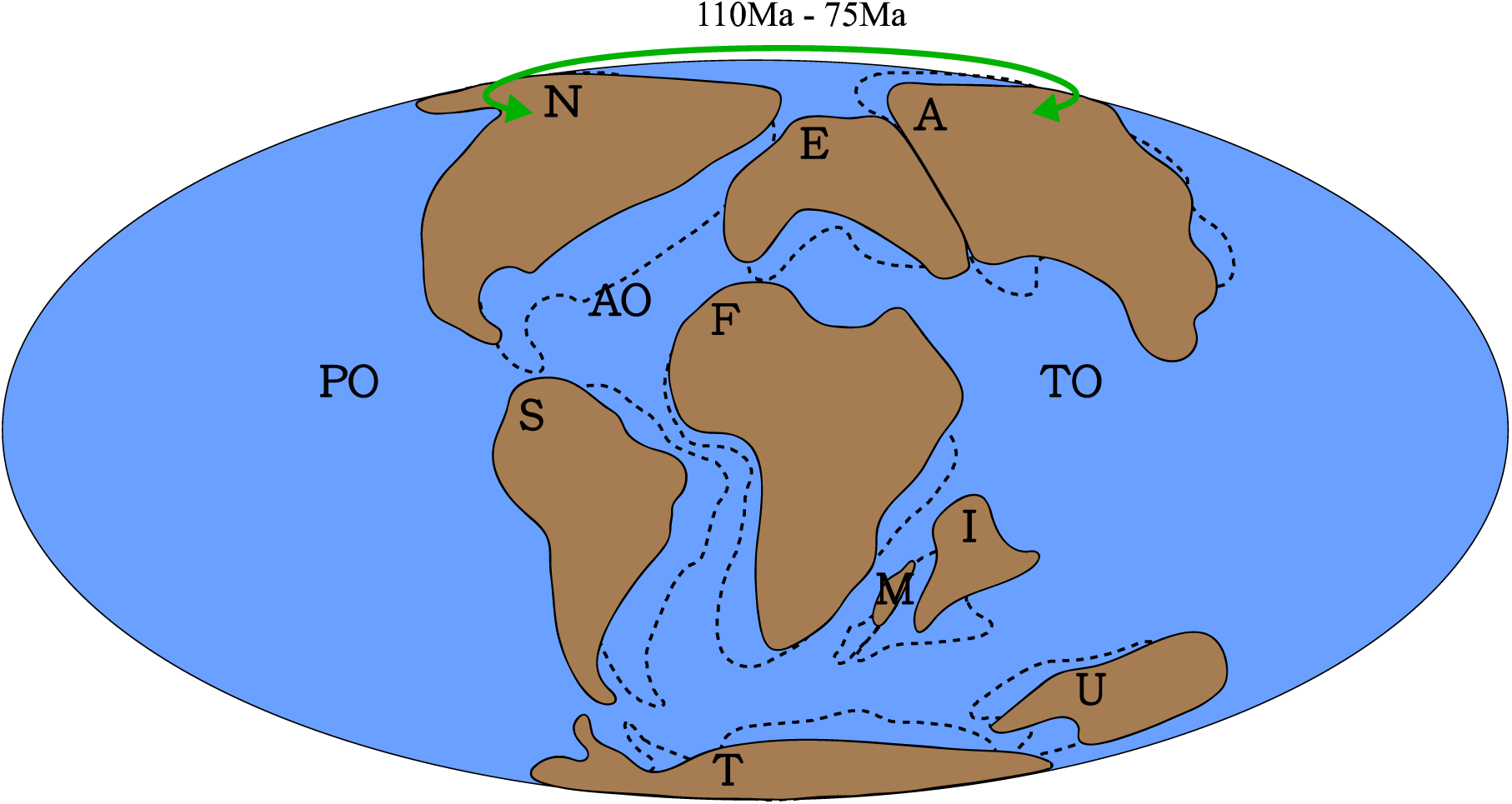
Hypothesis 4, Cretaceous North America-Asia faunal exchanges. This includes both the Early and Late Cretaceous establishments of the Bering land-bridge. The green arrowed line denotes the approximate dispersal directions and route. Dotted lines denote palaeogeography at 110Ma, solid lines denote palaeogeography at 75 Ma. Palaeomap after (Matthews, et al., 2016). Abbreviations: A, Asia; AO, Atlantic Ocean; E, Europe; F, Africa; I, India; M, Madagascar; N, North America; PO, Pacific Ocean; S, South America; T, Antarctica; TO, Tethys Ocean; U, Australia.

The dinosaurian biogeographic hypotheses we propose to test in this study include the existing hypotheses outlined above, as well as ones that we have modified or developed ourselves (Table 1). Existing hypotheses are mostly based on narrative or qualitative approaches, which has limited their accuracy as well as ability to undergo statistical testing. To address this issue, quantitative biogeographic techniques are examined below to identify the most suitable method to implement in this study.

**TABLE 1.**
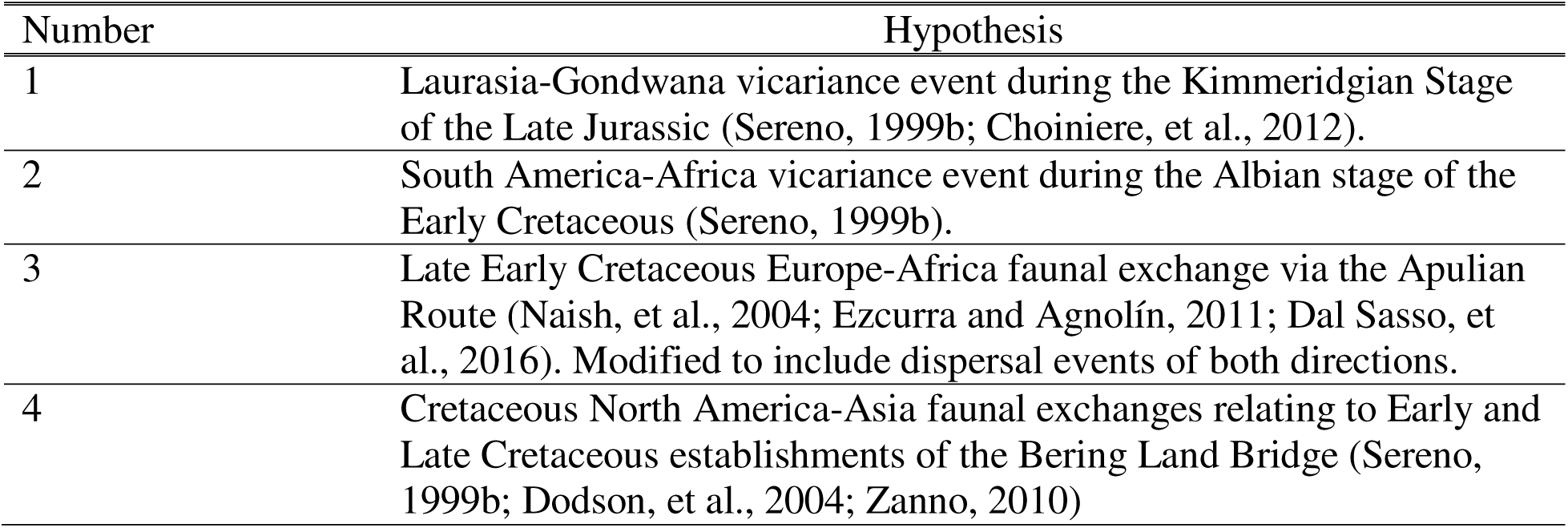
Biogeographic hypotheses tested in this study.

## Ancestral Crown Avian Biogeography

The Mesozoic biogeographic history of coelurosaurs set the stage for the early biogeographic history of crown birds. Although the stunning diversity of living birds and their easily observable nature would seem to simplify robust biogeographic inferences for the major clades of modern birds, deep crown bird biogeography has emerged as one of the most contentious issues in contemporary avian macroevolution (Cracraft and Claramunt, 2017; Mayr, 2017; Field and Hsiang, 2018).

Opposing views on crown bird historical biogeography relate to the observation that the early Cenozoic fossil record of crown birds frequently reveals unforeseen complexity in the distributions of major clades. For example, many major clades of extant birds are restricted to vestiges of Gondwana (South America, Africa and Australasia). As a result, quantitative ancestral biogeographic reconstructions invariably infer a Gondwanan origin of the avian crown group, and for many of the deepest nodes within the crown bird tree of life (Fig. 5) (Cracraft, 2001; Claramunt and Cracraft, 2015). However, the earliest known fossil stem group representatives of many such ‘Gondwanan’ clades derive from the Palaeogene of the Northern Hemisphere, entirely outside the modern geographic distributions of their crown clades, casting doubt on what the ancestral geographic distributions for these groups really were. This holds true for clades currently restricted to Africa such as Musophagiformes (Field and Hsiang, 2018) and Coliiformes (Houde and Olson, 1992; Mayr and Peters, 1998; Mayr, 2001; Zelenkov and Dyke, 2008; Ksepka and Clarke, 2009; Ksepka and Clarke, 2010a); Madagascar such as Leptosomiformes (Mayr, 2002a; 2002b; 2008); South America such as Cariamiformes (Mourer-Chauviré and Cheneval, 1983; Peters, 1995; Mourer-Chauviré, 1999; Mayr, 2000; 2002a; Mourer-Chauviré, 2006) and Opisthocomiformes (Mayr and De Pietri, 2014); and Australasia such as Podargiformes (Nesbitt, et al., 2011; Mayr, 2015). Indeed, the dynamic nature of crown bird biogeography is further evinced by the early Old World fossil record of clades presently restricted to the New World such as hummingbirds (Trochilidae) (Karhu, 1988; 1992; 1999; Mayr, 2003; 2004; Bochenski and Bochenski, 2008; Louchart, et al., 2008), and clades presently restricted to the Old World formerly distributed in the New World such as the roller + ground roller clade (Coracii) (Mayr, et al., 2004; Clarke, et al., 2009; Ksepka and Clarke, 2010b).

**Fig. 5.**
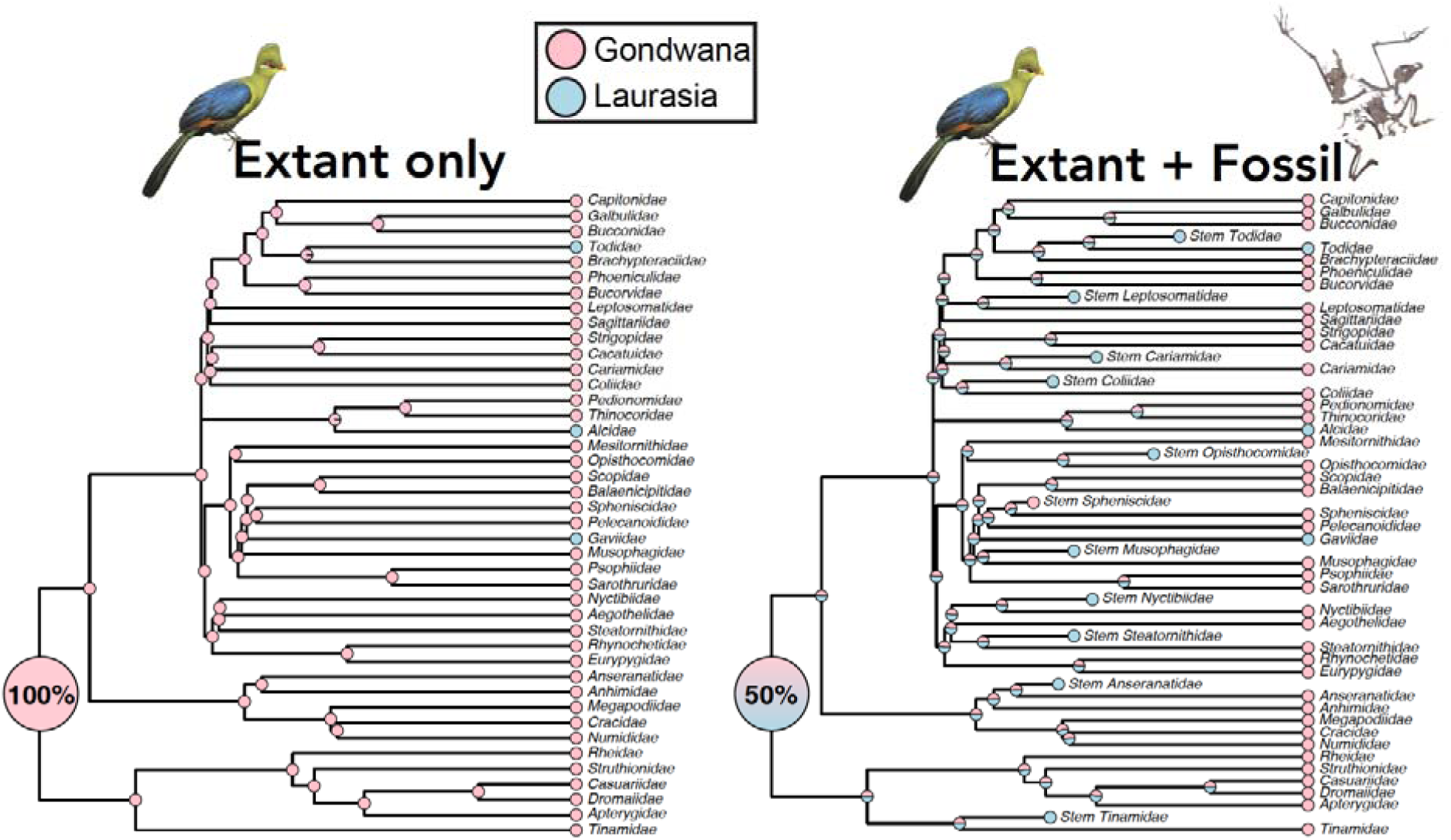
Illustration of conflict between ‘extant only’ biogeographic reconstructions for crown birds, and the crown bird fossil record (modified from (Field and Hsiang, 2018)). Extant only reconstructions infer a Gondwanan origin of crown birds with strong probability (**A**), whereas identical analyses incorporating the earliest fossil stem group representatives infer a markedly less robustly supported result (**B**).

Inclusion of early Cenozoic crown bird fossils in avian historical biogeographic reconstructions therefore has potential to erode confidence in erstwhile robust analytical reconstructions of crown bird historical biogeography (Fig. 5). Moreover, as the evolutionary timescale of crown birds has come into clearer focus (Feduccia, 2014; Jarvis, et al., 2014; Prum, et al., 2015; Berv and Field, 2018) (Field, et al., this volume), attempts to reconcile the ‘trans-Antarctic’ distributions of many groups of crown birds through Gondwanan vicariance (Cracraft, 2001) have emerged as untenable since Gondwanan breakup was largely complete by the time crown birds arose (Field and Hsiang, 2018). As a result, the true biogeographic origins of crown birds may be best regarded as uncertain at present; only future fossil discoveries of the earliest crown birds from the latest Cretaceous and earliest Cenozoic have potential to shed direct light on ancestral biogeographic distributions of crown birds (Field and Hsiang, 2018). As such, this article will not treat the biogeographic history of crown birds (the ‘living coelurosaurs’) and will instead focus solely on the Mesozoic biogeography of the major non-avian coelurosaur clades.

## Quantitative Biogeographic Methods

Quantitative biogeographic methods are mainly used for inferring the ancestral geographic distributions of species and clades, as well as the biogeographic processes that produced the observed species distribution. Quantitative analyses require phylogenetic trees of the target clades and some models or assumptions about the evolution of faunal distribution (Ronquist and Sanmartín, 2011). A statistical framework, including parsimony (to a wider extent) and likelihood, is used for formulating an analysis (Ronquist, 1997; Ree, 2005; Landis, et al., 2013; Matzke, 2013).

These historical biogeographic methods are analogous to phylogenetic analysis for inferring character evolution, in that the characters of the taxa are replaced with geographic distributions (Ree, 2005). Therefore, they share similar statistical frameworks, with parsimony, likelihood, and Bayesian methods all being applied in different quantitative biogeographic techniques (Ronquist, 1997; Ree, 2005; Landis, et al., 2013). Although the debate on the justification of different statistical frameworks in phylogenetic methods is heated (Goloboff, et al., 2018a; Goloboff, et al., 2018b; O’Reilly, et al., 2018), the selection of methodological approach is more straightforward and constrained than in historical biogeography. To date, no single method has yet been shown to have better performance in historical biogeographic analyses. Validation of the plethora of biogeographic techniques is beyond the scope of this project, though this work should be a priority in the future.

In most analytical approaches the whole geographic range of interest is divided into multiple smaller areas and taxa are assigned to one or more of these areas. Faunal distribution evolution models are simplified versions of biogeographic processes that operate on the defined geographical areas (Ronquist, 1997; Ree, 2005; Landis, et al., 2013; Matzke, 2013). For example, a dispersal event for a taxon corresponds to an increase in the number of distribution areas at a node and/or along a branch in a taxon phylogeny. Other distribution evolution models include regional extinction, sympatry, vicariance, and founder-event speciation (the latter occurring when one of the two daughter lineages of an ancestor acquires a different area to that ancestor, usually through dispersal across a barrier). Different quantitative biogeographic methods take different models into consideration, which will directly affect the results obtained (Matzke, 2013). Thus, a multi-model approach is recommended to more accurately identify biogeographical patterns.

The first widely used quantitative biogeographic analysis approach was Dispersal-Vicariance Analysis or DIVA (Ronquist, 1997), as implemented in the program FigTree. This parsimony-based approach utilises phylogenetic character optimisation methods and models dispersal, extinction and vicariance events. Each event is assigned with a cost (the cost of dispersal and regional extinction events are 1 whilst the cost of vicariance event is 0). The overall biogeographic history with the lowest cost is favoured (Ronquist, 1997). However, the time dimension is not considered in the analysis. Later, a likelihood framework was introduced in the form of the dispersal-extinction cladogenesis model (DEC) by assigning dispersal and extinction rates as free parameters, which can vary to give different overall likelihoods (Ree and Smith, 2008). Subset sympatry (one of the daughter lineages living in a subset of the ancestral range, while the other continues to occupy the whole ancestral range) and a limited form of vicariance (one of the daughter lineages occupies only one ancestral area, while the other occupies the rest of the ancestral range) are permitted, but widespread sympatry or vicariance are prohibited. By varying free parameter values, the ancestral state is reconstructed by maximising the overall likelihood of the whole biogeographic process with the branch lengths on phylogenetic trees i.e. evolutionary time is taken into consideration. Historical geographic changes can also be incorporated into the analysis using this method (Ree and Smith, 2008). Finally, in order to enhance the computational speed of biogeographic analysis, the program BayArea was developed with a Bayesian approach based on a likelihood framework, in which vicariance was prohibited (Landis, et al., 2013).

## Methodology

In this project, we adopt the R package, BioGeoBEARS (Matzke, 2013), for analyzing coelurosaurian biogeography. Likelihood versions of the biogeographic models in DIVA, DEC, and BayArea are incorporated in BioGeoBEARS, which allows the results of several different models to be more easily compared. DEC is the original Dispersal-Extinction Cladogenesis model (Ree and Smith, 2008). DIVALIKE is a likelihood version of Dispersal-Vicariance Analysis (Ronquist, 1997). Unlike DEC, the DIVALIKE model disallows subset sympatry, but permits widespread vicariance (i.e. two daughter lineages dividing up the ancestral range and both sharing more than one area). BAYAREALIKE is a likelihood-based version of the BayArea program (Landis, et al., 2013). In BAYAREALIKE, the two daughter lineages of a given ancestor inherit the same area distribution as that ancestor. As a consequence, the BAYAREALIKE model allows widespread sympatry (i.e. for any ancestral range occupying more than one area, a daughter lineage copies it), which is prohibited in both the DEC and DIVALIKE models. Like BayArea, BAYAREALIKE also disallows vicariance events. All three models assume narrow sympatry i.e. spontaneous range copying of single area ancestral ranges and set dispersal and regional extinction rate as free parameters. In BioGeoBEARS, founder-events are included as a separate range switching process (termed the J parameter), which can be considered as a rapid dispersal event. This creates three new variants of the three models, giving a total of 6 possible model comparisons per biogeographic dataset: DEC, DIVALIKE, BAYAREALIKE, DEC+J, DIVALIKE+J, and BAYAREALIKE+J (Matzke, 2013). Standard statistical comparison with LnL or the Akaike Information Criterion (AIC) can be performed to identify the model(s) that best fits the data.

Applying BioGeoBEARS has three notable advantages over existing methods: 1) different models are compared based on the same dataset (as mentioned above); 2) geological time spans are considered in calculating the likelihood of faunal distribution evolution; 3) palaeogeographic constraints can be implemented to inform the analysis about area connectedness during the Mesozoic. At this stage, these three features cannot be achieved by any current software based on a parsimony framework or Bayesian approach.

BioGeoBEARS considers time in calculating the likelihood of faunal distribution evolution. All things being equal, a biogeographic event should have a higher probability of occurring over a longer period of time. This time span, which is the branch length of each lineage, is taken into consideration within the likelihood framework through the relationship:

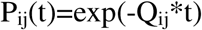

Where P is the event likelihood, Q is the universal event rate (dispersal, local extinction, vicariance, founder-events) and t is the branch length (Matzke, 2013). Although assigning one universal rate value to each biogeographic event is still over simplistic, the consideration of time, which is not achieved within a parsimony framework yet, remains an important advantage of analyses currently.

BioGeoBEARS allows palaeogeographic constraints with known temporal ranges to be incorporated into an analysis (Matzke, 2013). This ensures that continents which are connected and separated are assigned different dispersal probabilities despite one universal dispersal rate. To do so, dispersal multipliers are introduced, whose product with the universal dispersal rate will be the new regional dispersal rate used to calculate the likelihood of the dispersal event in question. The dispersal multipliers for connected geographic ranges are set to 1 while those for separated ranges are set to 0.000001 (a low value does not rule out such dispersal events but implies that they are highly unlikely). Here, we follow the protocol of Poropat, Mannion et al. (2016), when dealing with regions that are separated by shallow seas or uncertain geographic barriers, the value 0.5 is assigned to the dispersal multiplier between those regions, which act as our starting (or normal) constraints (see Poropat, et al. (2016: for further details).

Within each analysis, the overall likelihood of reproducing the dataset given the model is computed, and the overall biogeographic process with the maximum probability (likelihood) will be the best fit result. The Akaike information criterion (AIC) and natural log of the process likelihood (LnL) will be calculated to infer the quality of data fit. A smaller AIC and a larger (less negative) LnL indicate a better fit of the data given the model tested.

We divide land areas on the Earth’s surface into 8 geographic units, namely Africa, Asia, Australia, Europe, India, Madagascar, North America and South America (although there are no taxa from India or Australia in our dataset). Each taxon is assigned to the areas according to data reviewed in the section *Geographic and Temporal Distributions of Coelurosaurs.* These data were obtained by referring back to holotype descriptions and other literature [there should be a table or appendix that gives the geographic ranges and temporal ranges for each terminal taxon, with supporting references, and this should be referred to here We will add this as a supplemental URL link – these is no space to print the appendix in the volume.

The coelurosaur phylogenetic tree with time calibration applied in the analysis is shown in Figure 1. This is a maximum agreement subtree of a recent analysis of the Theropod Working Group (TWiG) phylogenetic matrix (see Brusatte, et al. (2014: for further details). The palaeo-geographic constraints are those used by Xu, et al. (2018:, as modified from those summarised by Poropat et al. (2016) by better constraining the opening and closing of the Russian Platform sea between Asia and Europe during the Jurassic (Xu, et al., 2018). They are represented by 23 dispersal multiplier matrices corresponding to 23 time-slices from the Middle Jurassic to the Late Cretaceous. Four analyses with starting, relaxed, harsh as well as no palaeogeographic constraints were carried out. Relaxed constraints set all 0.5 dispersal multiplier values to 1, harsh constraints set all 0.5 dispersal multipliers to 0.000001. These analyses were repeated for all six biogeographic models giving a total of 24 comparisons.

## RESULTS

The relative fit of the 24 analyses to the data are summarized in Table 2, with the corresponding parameter values listed in Table 3.

**TABLE 2.**
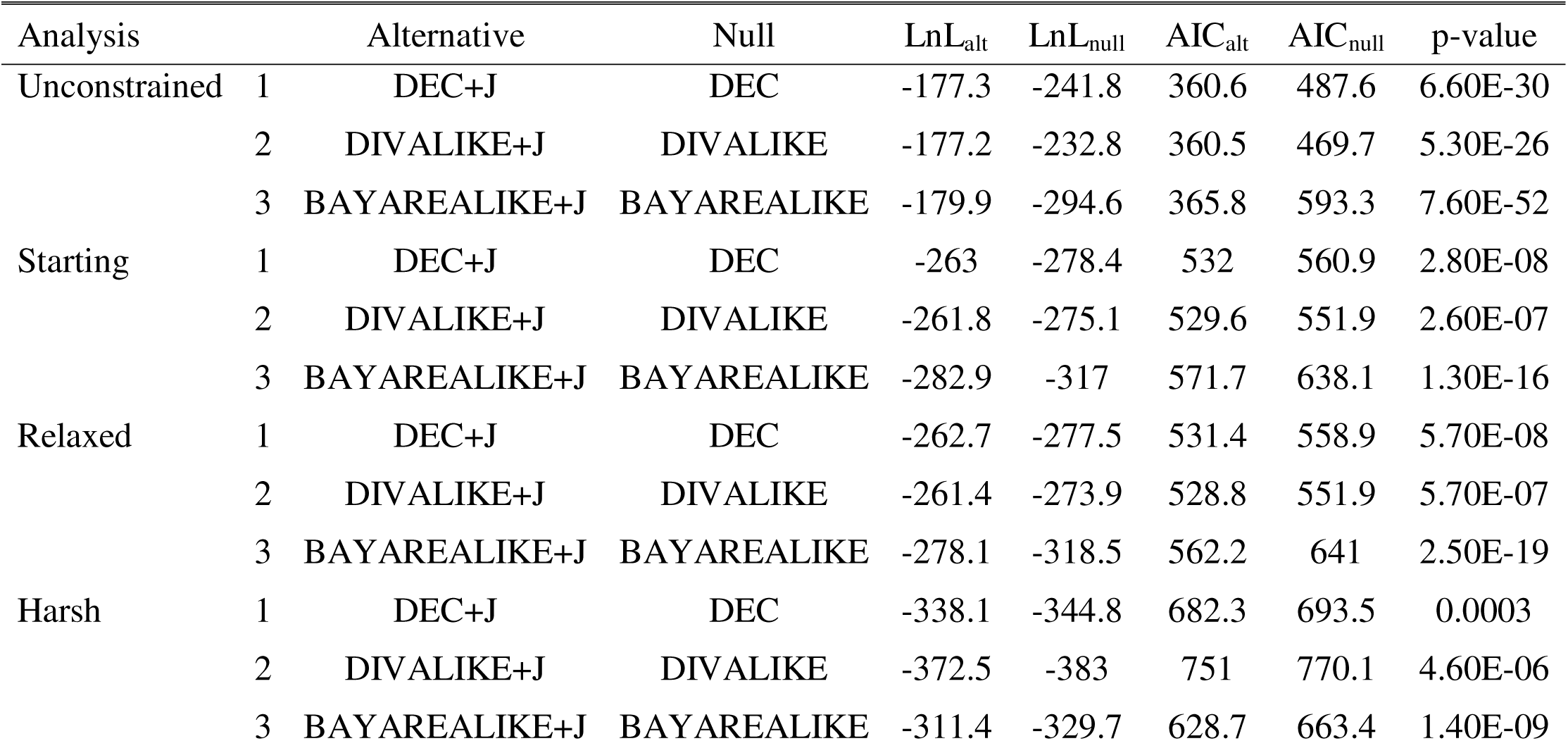
Relative performance of the six biogeographic models under the unconstrained and three palaeogeographically constraint conditions. Abbreviations: Alt, alternative model; null, null model; LnL, natural log of the process likelihood; p-value, p-value of the Likelihood Ratio Test; AIC, Akaike Information Criterion.

**TABLE 3.** Parameter values for the six biogeographic models under the unconstrained and three palaeogeographically constraint conditions. Abbreviations: LnL, natural log of the process likelihood; d, dispersal rate; e, regional extinction rate; j, founder-event rate; AIC, Akaike Information Criterion; AICweight, Akaike Information Criterion weight.

|  | Model | LnL | d | e | j | AIC | AICweight |
| --- | --- | --- | --- | --- | --- | --- | --- |
| Unconstrained | DEC | -241.8 | 0.001 | 0.0021 | 0 | 487.6 | 1.20E-28 |
|  | DEC+J | -177.3 | 1.00E-12 | 1.00E-12 | 0.023 | 360.6 | 0.47 |
|  | DIVALIKE | -232.8 | 0.0012 | 2.00E-09 | 0 | 469.7 | 9.50E-25 |
|  | DIVALIKE+J | -177.2 | 1.00E-12 | 1.00E-12 | 0.024 | 360.5 | 0.49 |
|  | BAYAREALIKE | -294.6 | 0.0011 | 0.015 | 0 | 593.3 | 1.40E-51 |
|  | BAYAREALIKE+J | -179.9 | 1.00E-12 | 1.00E-12 | 0.024 | 365.8 | 0.035 |
| Starting | DEC | -278.4 | 0.014 | 0.0084 | 0 | 560.9 | 1.30E-07 |
|  | DEC+J | -263 | 0.0083 | 0.0059 | 0.1 | 532 | 0.23 |
|  | DIVALIKE | -275.1 | 0.016 | 0.0077 | 0 | 554.1 | 3.70E-06 |
|  | DIVALIKE+J | -261.8 | 0.0099 | 0.0062 | 0.1 | 529.6 | 0.77 |
|  | BAYAREALIKE | -317 | 0.013 | 0.022 | 0 | 638.1 | 2.10E-24 |
|  | BAYAREALIKE+J | -282.9 | 0.0013 | 0.015 | 0.051 | 571.7 | 5.50E-10 |
| Relaxed | DEC | -277.5 | 0.011 | 0.0081 | 0 | 558.9 | 2.30E-07 |
|  | DEC+J | -262.7 | 0.0068 | 0.0059 | 0.083 | 531.4 | 0.21 |
|  | DIVALIKE | -273.9 | 0.013 | 0.0075 | 0 | 551.9 | 7.90E-06 |
|  | DIVALIKE+J | -261.4 | 0.0083 | 0.0062 | 0.082 | 528.8 | 0.79 |
|  | BAYAREALIKE | -318.5 | 0.011 | 0.022 | 0 | 641 | 3.50E-25 |
|  | BAYAREALIKE+J | -278.1 | 0.0063 | 0.0082 | 0.12 | 562.2 | 4.50E-08 |
| Harsh | DEC | -344.8 | 0.021 | 0.0086 | 0 | 693.5 | 8.60E-15 |
|  | DEC+J | -338.1 | 0.015 | 0.0072 | 0.082 | 682.3 | 2.40E-12 |
|  | DIVALIKE | -383 | 0.023 | 0.008 | 0 | 770.1 | 2.00E-31 |
|  | DIVALIKE+J | -372.5 | 0.015 | 0.0065 | 0.1 | 751 | 2.70E-27 |
|  | BAYAREALIKE | -329.7 | 0.0056 | 0.024 | 0 | 663.4 | 2.90E-08 |
|  | BAYAREALIKE+J | -311.4 | 0.0037 | 0.019 | 0.049 | 628.7 | 1 |

As shown in Table 2, the AIC and LnL values indicate that the unconstrained analysis, the analysis with the starting constraints, and the analysis with the relaxed constraints prefer the DIVALIKE+J model, while the analysis with the harsh constraints prefers the BAYAREALIKE+J model.

It is expected that the unconstrained analysis will typically perform better than the constrained ones for three reasons. First, without geographic constraints, the dispersal rate remains constant between any two geographic areas throughout coelurosaurian evolutionary history. Under these conditions, unlikely dispersal events will not be prohibited and will have the same probability of occurring as palaeogeographically more plausible dispersals. For example, Asian taxa may be able to directly disperse to South America, which to a large extent simplifies the biogeographic processes. As a result, the overall likelihood of the unconstrained processes should be expected to be higher than the constrained ones’ as a result of over-simplification of the biogeographic processes. Second, vicariance events can take place whenever a fauna is distributed across more than one geographic area. Since the overall dispersal rate is low between continents, any fauna having cross-continent biogeographic ranges is separated by ‘geographic barriers’ as understood by the statistical framework. Due to the high flexibility for vicariance events in an unconstrained analysis, low probability regional extinction events will frequently be replaced by “must-happen” vicariance, which boosts the overall process likelihood (in models that allow vicariance, the vicariance rate is set to be close to 1). Third, founder-events, as a fast version of dispersal, will be more frequently applied in an unconstrained analysis. A founder-event is similar to instant range switching of an ancestral state, which shares the same rate as a dispersal event. In an unconstrained analysis, it not only mimics the process of dispersal, but also does not need low probability regional extinction events to account for taxa that occupy just a single area. Therefore, founder-events can accommodate any taxon that occupies an area that differs from its ancestral state, and this will be more statistically favourable than the estimation of vicariance events to explain the observed distributions. The large LnL contrasts between models with (+J models) and without founder-event speciation are also expected because of the statistical preference for models that include this parameter. The predicted high likelihoods achieved by of the unconstrained analyses is confirmed in Table 2 where the LnL values for all six biogeographic models are less negative than those in any constrained analyses. Thus, in spite of the low AIC and high LnL values, the results of the unconstrained analysis should be treated with caution because they do not consider palaeogeographic constraints. The absence of such information relevant to the direction and probability of faunal dispersal is ultimately likely to lead to less accurate estimations of dispersal events e.g. direct dispersal events from Asia to South America across the Mesozoic Pacific Ocean.

Within the constrained analyses, the starting constraints and relaxed constraints give results that agree with each other on the most preferred model, DIVALIKE+J, while the harsh constraints suggest a preference for BAYAREALIKE+J. Here, we accept DIVALIKE+J as the best supported model based on the following reasons:

1. DIVALIKE+J is preferred in the analysis with the starting constraints, which to our knowledge best reflects Mesozoic geography. Such model preference is also supported in the analysis with the relaxed constraints.
2. The ancestral state reconstruction of the DIVALIKE+J model provides realistic situations and processes. This is discussed in more detail below.
3. The results of the BAYAREALIKE+J model (including the one under the harsh geographic constraints) estimate several occurrences of ancestors that were present solely in South America and Asia during the Cretaceous. Such ancestral area estimations are not realistic since the two continents were separated by large oceans during that time (Scotese, 2001) and it is highly unlikely that faunal exchange ever happened. Events of this type are frequent in the BAYAREALIKE+J model, because a high value (nearly 1) is assigned to the widespread sympatry process rate. Some authors (Poropat, et al., 2016; Xu, et al., 2018) interpret such unlikely ancestral distributions as resulting from the impacts of uneven sampling of the fossil record. In reality, such widespread sympatry cannot happen without faunal exchange across these areas, but this is neglected in the model. The unrealistic results and over-simplified biogeographic processes justify the rejection of this model.
4. The Harsh biogeographic constraints might not accurately reflect palaeogeography and the true dispersal capabilities of coelurosaurs. This is because they treat all uncertain connections and shallow seas as geographic barriers, which largely isolates the individual continents throughout the Cretaceous. The harsh constraints could therefore be under-estimating the dispersal ability of coelurosaurs. Analogous to the modern Madagascar fauna (Ali and Huber, 2010), coelurosaurs - particularly small ones with or without aerodynamic capabilities - might have been capable of crossing relatively narrow channels or shallow water ways over geological time scales.

In the various constrained analyses, the +J models still perform better than non+J models, although the differences are not as large as in the unconstrained analysis (as suggested by p-values in Table 2). This phenomenon is also because founder-events as alternative dispersal processes are statistically more favourable than dispersal events. However, with the implementation of palaeogeographic constraints, this flexibility is to some extent restricted and vicariance events are preferred if no feasible dispersal route is allowed, thereby reducing the likelihood gap of +J models and corresponding non+J models. However, which particular events happened during coelurosaurian evolution, whether founder-events or dispersals, cannot be determined by these analyses.

In the analyses with the starting and relaxed constraints, the DIVALIKE+J and the DEC+J models perform nearly equally well with close AIC and LnL values. Both of the models allow narrow vicariance and founder-events, while DIVALIKE+J allows widespread vicariance but DEC+J allows subset sympatry. Due to the fact that nearly all taxa in our analysis only occupy single geographic areas and large-scale connections of these units were absent after the breakup of Pangaea, biogeographic processes that result in widespread ancestral species probably did not play a major role in coelurosaurian evolution. This might explain the similar results of the DIVALIKE+J and DEC+J models. The ancestral area estimations of the two models also provide similar information of biogeographic history. Because the DIVALIKE+J model is slightly favoured in terms of AIC and LnL value, we use its results in our discussion of coelurosaurian biogeography. The results of the starting constraints are used as the basis of our discussion here because they represent the most conservative palaeogeography among the three constraints (starting, relaxed, harsh).

The major biogeographic processes during coelurosaurian evolution include inter-continental dispersals, regional extinctions, continent-scale vicariance events, and continental scale founder-events, as our area units are at the continental scale our results cannot capture intra-continental and island-scale biogeographic patterns.

Important coelurosaurian dispersal pathways appear to include the earliest Cretaceous Apulian Route connecting southwestern Europe and north-western Africa and the Bering Land Bridge connecting north-eastern Asia and north-western North America during the late Early Cretaceous and latest Cretaceous. The ancestral area estimation suggests that the Apulian Route (Fig. 3) played an important role in shaping coelurosaurian biogeography. It appears to account for the South American occurrence of the Early Cretaceous compsognathid *Mirischia* and the Late Cretaceous avians *Patagopteryx* and *Neuquenornis*, while their closest relatives were present in Asia (although multiple inter-continental dispersals probably took place). Thus, our results support Europe-Africa faunal exchange as proposed in Hypothesis 3 (Fig. 3, Table 1). The establishment of the Bering Land Bridge enabled direct Asia-North America dispersal without a transit via Europe. This dispersal route is the most frequently used by coelurosaurs as inferred from our results. Single dispersal events during the first establishment of the Bering Land Bridge in the late Early Cretaceous (Fig. 3) potentially explain the Asian occurrence of the dromaeosaurid *Achillobator*, the troodontid *Troodon*, the dromaeosaurid *Dakotaptor*, as well as the Asian occurrence of the dromaeosaurid *Achillobator*. Our results therefore partially support Hypothesis 4 (Fig. 4, Table 1). Furthermore, the North American occurrence of the alvarezsaurid *Albertonykus* and ornithomimids *Struthiomimus* and *Ornithomimus* as well as the Asian occurrence of the tyrannosaurid *Alioramus*, can be attributed to ancestral dispersal via the Bering Land Bridge during its second connection in the latest Cretaceous (Fig. 4). Thus, together, our results support Hypothesis 4 in full (Table 1), indicating that the two episodes of connection across the Bering Strait probably facilitated important faunal exchanges that produced most of the Asia-North America coelurosaur occurrences throughout the Cretaceous.

While dispersal and extinction are part of all six BioGeoBEARS models, the support for the DIVALIKE+J model in particular indicates important roles for vicariance and founder-event speciation in coelurosaurian evolution. In contrast, the BAYAREALIKE models, which do not allow vicariance, are not supported except when particularly stringent (harsh) palaeogeographic constraints are imposed. Such vicariance events can be recognized in the ancestral area estimations and potentially linked to continental disconnection events during the Mesozoic. This includes Gondwana-Laurasia vicariance resulting from the breakup of Pangaea in the Middle Jurassic, which explains the occurrence of the most stemward coelurosaur *Bicentenaria* in South America and other stemward taxa in Laurasia. Thus, our results support Hypothesis 1 (Fig. 1, Table 1). We propose an Asia-North America vicariance event due to the breakdown of the first Bering Land Bridge during the early Late Cretaceous (Hypothesis 5), which could explain the occurrence of *Nothronychus* in North America and *Nanshiungosaurus* in Asia based on our results (Figure 6). Such vicariance events might have been frequent and repeated because of the intermittent breakdown and reformation of fine-scale geographic connections (such as carbonate platforms, land-bridges across shallow epicontinental seas etc.) between different continents after the breakup of Pangaea. The South America-Africa vicariance event (Hypothesis 2) is not recognised in our results, most probably because of the dearth of African coelurosaurs in the late Early Cretaceous fossil record. Within the DIVALIKE and DEC models, those with founder-events (i.e. +J) perform better than those without, which implies that founder-events may have also frequently occurred during coelurosaurian evolution. However, on a continental scale, founder-events at ancestral nodes, although statistically preferred, become similar to within-lineage dispersal events from a biological point of view, since at least one faunal dispersal event is necessary. Hence, no absolute or relative frequencies of founder-events can be inferred from our results.

**Fig. 6.**
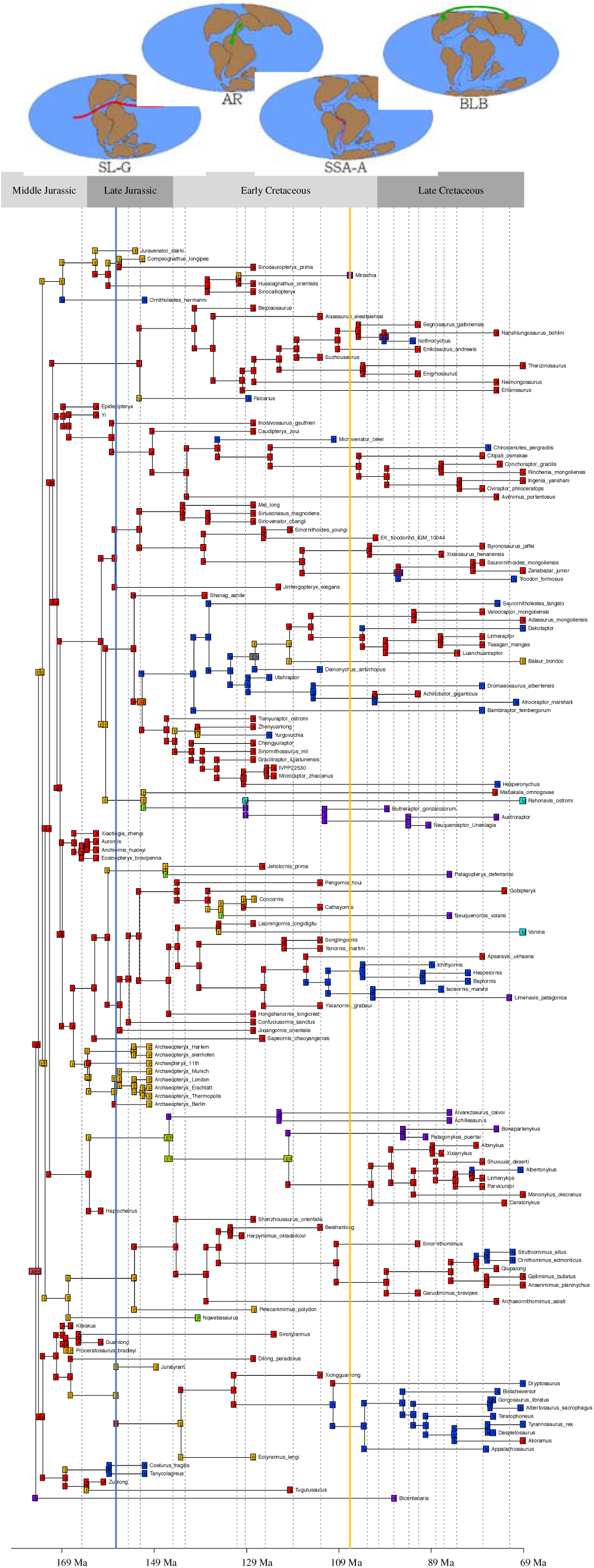
Ancestral area estimation applying the DIVALIKE+J model with starting constraints to a dated coelurosaurian phylogeny. The different geographic areas are denoted by letters as follows: A: Asia; E: Europe; F: Africa; M: Madagascar; N: North America; S: South America. Green shading denotes the period when the Apulian Route (AR) connected northeast Africa and southwest Europe, whilst red shadings denote Bering Land-bridge (BLB) connections. The blue line denotes the time of separation between Laurasia and Gondwana (SL-G), whilst the yellow line denotes the time of separation between South America and Africa (SSA-A).

Europe appears to have played a key role as a dispersal centre and gateway during coelurosaurian evolution. Although only having a few coelurosaur occurrences, Europe forms all or part of the geographic range for many ancestral nodes in our results (Fig. 7). On the one hand, some coelurosaurian faunas might have originated in Europe and dispersed to adjacent continents including Asia, North America and Africa. Examples in our results include the inferred origin of Compsognathidae in Europe, followed by European-centred dispersal events to Asia and North America, and the inferred origin of Ornithomimosauria in Europe, followed by dispersals to Asia and Africa. Moreover, some coelurosaurian faunas might have experienced long-distance dispersal via Europe. Before the late Early Cretaceous, when the eastern part of Asia and western part of North America were still separated from each other directly, faunal exchange between these two continents required travel via Europe. The ancestor of the dromaeosaurid *Yurgovuchia*, most likely took this route. As mentioned above, Laurasian-Gondwanan faunal exchange was only possible via the Apulian Route connecting Europe and Africa, so crownward coelurosaurs from Gondwanan landmasses should have migrated from Laurasia via Europe. Therefore, Europe should have been an essential geographic area for shaping coelurosaurian biogeography as both a dispersal centre and a geographical gateway (Fig. 8).

**Fig. 7.**
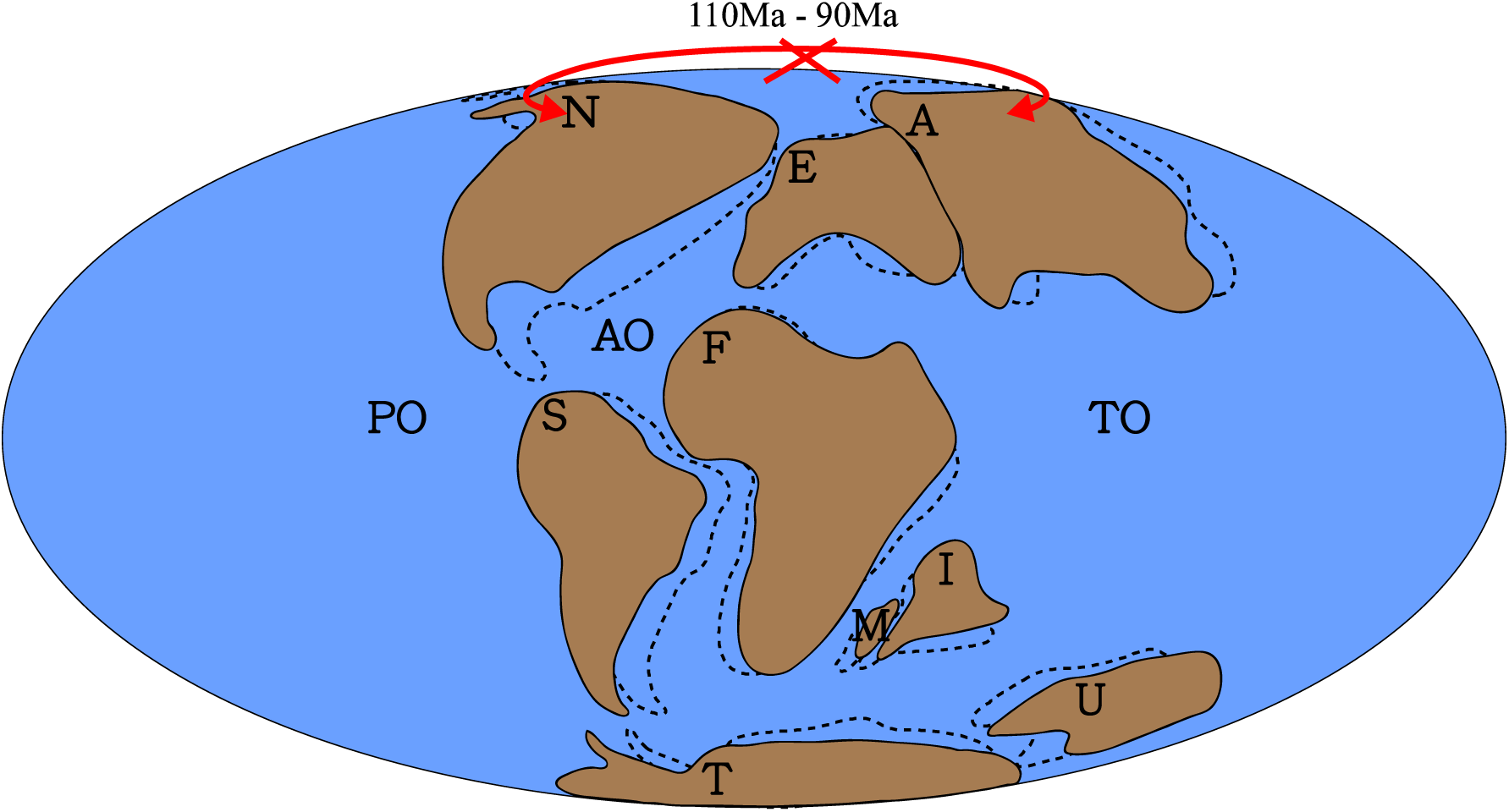
Hypothesis 5, North America-Asia vicariance event caused by the breakdown of the Bering Land-bridge during the early Late Cretaceous. The marine barrier between northeast Asia and northwest North America was re-established in the Cenomanian Stage. The red cross denotes the approximate position of the hypothesized biogeographical barrier (Bering Strait); Dotted lines denote palaeogeography at 110 Ma, whilst solid lines denote it at 90 Ma. Palaeomap after (Matthews, et al., 2016). Abbreviations: A, Asia; AO, Atlantic Ocean; E, Europe; F, Africa; I, India; M, Madagascar; N, North America; PO, Pacific Ocean; S, South America; T, Antarctica; TO, Tethys Ocean; U, Australia.

**Fig. 8.**
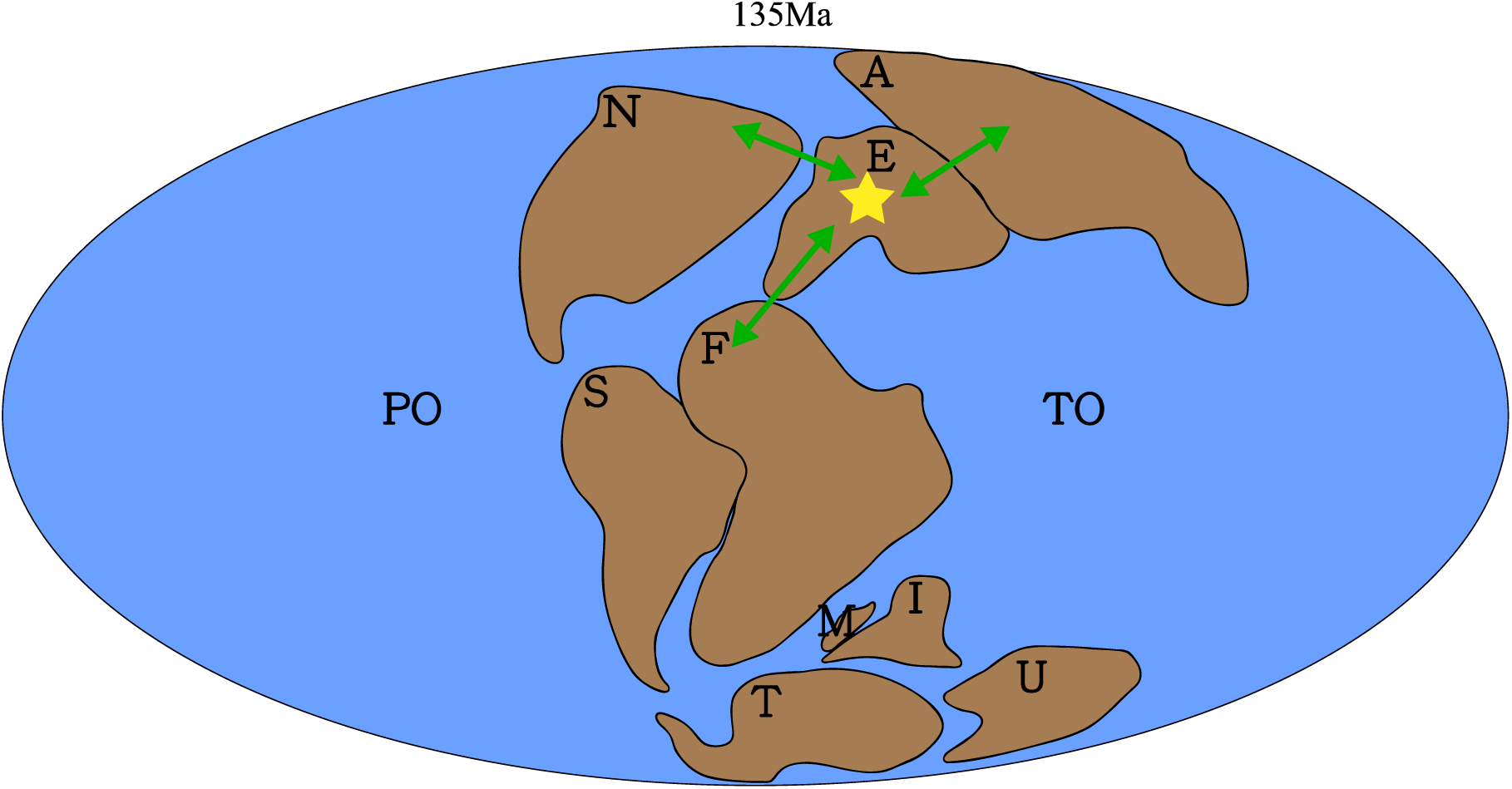
Europe as a dispersal center and geographical gateway, especially during the Early Cretaceous. The green arrowed lines denote possible dispersal directions and approximate dispersal routes; Solid lines denote palaeogeography at 135 Ma. Palaeomap after (Matthews, et al., 2016). Abbreviations: A, Asia; E, Europe; F, Africa; I, India; M, Madagascar; N, North America; PO, Pacific Ocean; S, South America; T, Antarctica; TO, Tethys Ocean; U, Australia.

## DISCUSSION

Our results confirm that continent-scale vicariance was probably an important biogeographic process influencing coelurosaurian evolution. This is consistent with many qualitative assessments in the literature (Sereno, 1999b; Fastovsky and Weishampel, 2005; Makovicky, et al., 2005; Choiniere, et al., 2012), and also agrees with the quantitative analysis of dinosaur biogeography by Upchurch *et al*. (2002b). Most workers recognised the importance of continental fragmentation in producing geographic barriers during the late Mesozoic, including the separation of Laurasia and Gondwana, the opening of the north Atlantic and the isolation of Gondwanan landmasses. Such vicariance events are seen in our results: for example, vicariance induced by the Middle Jurassic breakup of Pangaea led to the occurrence of the stem coelurosaur *Bicentenaria* in South America and other stemward taxa in Laurasia (Hypothesis 1). Similar vicariance patterns are also recognized within other terrestrial animal faunas at that time, such as dryolestoid mammals and eilenodontine sphenodontians (Makovicky, et al., 2005). However, we also note additional vicariance events that have not figured prominently in previous studies. An example of the latter is the apparent impact of the destruction of Land-bridges such as that across the Bering Strait (Hypothesis 5). Ephemeral land-bridges that reconnect separate areas, after continental fragmentation, were established from time to time throughout the Cretaceous, enabling inter-continental faunal dispersals. After the loss of these land bridges, the continents became isolated from each other once again and dispersed populations were separated by an oceanic barrier, which eventually caused vicariance. Such vicariance events are observed in the Therizinosauroidea and Troodontidae after the disappearance of the Bering Land Bridge during the early Late Cretaceous, and also within Alvarezsauroidea after the loss of the Apulian Route during the mid-Early Cretaceous.

Besides vicariance, the impact of other biogeographic processes on coelurosaurian evolution can also be recognized in our results. As suggested by multiple authors (Sereno, 1999b; Brusatte, et al., 2013; Dunhill, et al., 2016), dispersal played a major role in shaping coelurosaurian biogeography. The results again confirm an important role for the Bering Land Bridge (Sereno, 1999b) (Hypothesis 4) and Turgai Sea (Jin, et al., 2012; Allain, et al., 2014). However, we also support the Apulian Route as important for coelurosaurian faunal exchange (Hypothesis 3), an idea that has been somewhat neglected in previous studies. The formation of the Apulian Route in the late Early Cretaceous marked the first Laurasia-Gondwana connection after the breakup of Pangaea during the Middle Jurassic. Other taxa are believed to have crossed from Gondwana to Laurasia via this route in the Cretaceous, including carcharodontosaurids (Brusatte, et al., 2009), and rebbachisaurid (Sereno, et al., 2007) and titanosaurian sauropods (Dal Sasso, et al., 2016). If intermediate coelurosaurian taxa from Gondwana are not derived from vicariance induced by Pangaean breakup, they most probably arrived from Laurasian landmasses. Although such ‘Laurasian arrival’ hypotheses have been suggested (Naish, et al., 2004; Foth and Rauhut, 2017), these biogeographic events were not linked by these authors to the only feasible Laurasia-Gondwana dispersal route known in the Early Cretaceous. In our results, many inferred dispersal events via the Apulian Route explain South American coelurosaurian occurrences of their Laurasian relatives, but discoveries of closely-related African taxa are lacking. Therefore, new African coelurosaur discoveries will be crucial for testing Apulian Route dispersal events.

Related to the impacts of the Apulian Route and trans-Turgai land bridges, our analyses indicate that Europe might have been both a dispersal centre and a geographical gateway in coelurosaurian evolution, especially before the Barremian. As demonstrated in the results section, several lineages are inferred to have had stemward forms in Europe and then dispersed to other continents, while faunas from other lineages disperse from one continent to another via Europe. In the former case, ancestral faunas might have migrated from Europe to North America before the full establishment of the north Atlantic (e.g. stem coelurosaurs like *Ornitholestes* and the Compsognathidae). They might also have migrated from Europe to Asia when terrestrial routes existed across the Turgai Sea (e.g. Compsognathidae as evidenced by the ancestral area estimations for the *Sinosauropteryx* lineage), and to Africa via the Apulian Route during the Early Cretaceous (e.g. Ornithomimosauria as evident from *Nqwebasaurus*), and from there onto other Gondwanan landmasses. Even without these Cretaceous land bridges, Europe is likely to have played a central role in coelurosaurian dispersal. In particular, ancestral faunas might have migrated between Asia and North America via Europe (e.g. Asia to North America dispersal in *Coelurus*), and from Laurasia to Gondwanan landmasses via Europe before the breakup of Gondwana (e.g. Asia to South America from Asia in *Alvarezsaurus*, consistent with the results of Xu, et al. (2018) on alvarezsauroid biogeography). This conclusion is consistent with the result found by Dunhill et al. (2016) that European dinosaurs had strong direct connections with those in adjacent continents during the Jurassic and Early Cretaceous. They also showed a high degree of connectivity between North America and Asia, Asia and Africa, Africa and North America during the Early Cretaceous (Dunhill, et al., 2016), which can be explained by faunal exchange events via Europe as a dispersal gateway. For example, Laurasian continents shared a similar ankylosaurian (Ősi, 2005), hadrosauroid (Prieto-Marquez, et al., 2006; Dalla Vecchia, 2009), non-dinosaurian archosaur (Ezcurra and Agnolín, 2012) and gobiconodontid mammal (Cuenca-Bescós and Canudo, 2003) faunas, implying faunal exchange between Europe, Asia and North America. There were also European and Gondwanan faunas that were closely related (Dalla Vecchia, 2003; Gheerbrant and Rage, 2006), including spinosaurid theropods (Charig and Milner, 1997; Ruiz-Omeñaca, et al., 2005), sauropods (Canudo, et al., 2008), and thereuodontid mammals (Kielan-Jaworowska, et al., 2004), which can be attributed to faunal exchanges via the Apulian Route. Some Pangaean faunas, including spinosaurid theropods from Asia (Buffetaut, et al., 2008), titanosaurian sauropods from Asia and North America (Salgado, et al., 1997; Wilson, 2002), and crocodyliforms from Asia (Wu and Sues, 1996), had their closest relatives in Gondwana, and probably arrived from southern continents via Europe. Thus, the events estimated here for European coelurosaurs and their relatives elsewhere, are probably part of a widespread pattern imposed on multiple terrestrial clades by Pangaean fragmentation and the subsequent creation and destruction of key land bridges.

Finally, our results also imply that regional extinction played an important role in coelurosaurian evolution, as suggested by Sereno (1999b). Given frequent dispersals, continental-scale extinction events are necessary to account for the single-area unit-distributions of most coelurosaurian taxa. That regional extinction and dispersal were of comparable importance throughout coelurosaurian evolution is indicated by the similar values of the extinction and dispersal rates obtained in the analyses (Table 3).

## CONCLUSIONS

There are several uncertainties in our analyses that we should bear in mind. The major ones include: the accuracy of palaeogeographic reconstructions; errors in phylogenetic tree topology and node dating; spatiotemporal sampling biases; and our lack of definitive knowledge of the dispersal abilities of different coelurosaurian clades. Although the phylogenetic tree used here is better sampled than any examined in previous analyses of coelurosaurian biogeography, no single tree includes all known taxa. Phylogenies also change as new morphological data become available, which could alter the sequence of inferred biogeographic processes. New fossil discoveries from continents with rare or previously unknown records of particular clades are likely to modify our conclusions in the future (e.g. by changing tree topology, taxon ranges, origination dates and so on, which in turn could favour other biogeographic patterns and processes). The accuracy of our results is also affected by as yet unquantified preservation biases, which will distort the observed spatial and temporal distribution of coelurosaurs (Upchurch, et al., 2011). The second major uncertainty is regarding coelurosaurian dispersal ability. Coelurosaurian body sizes vary by six orders of magnitude, so their dispersal probability across narrow seaways probably varies significantly. Avians and probably some non-avian paravians developed powered flight ability, which may have freed them from the constraints of conventional terrestrial dispersal corridors. However, these abilities are still not well understood. Future analyses will need to better quantify dispersal ability if we are to move away from the uniform dispersal probabilities assumed in this study, although Mesozoic avians do not appear to have had the ability to cross expansive oceans using their own flight (Allen, et al., 2013). One group of comparable flying vertebrates is pterosaurs. Although this clade had different biogeographic patterns from other Mesozoic terrestrial vertebrates, quantitative biogeographic analysis revealed little vicariance and dispersal signal, but high levels of sympatry, which indicates rare cross-ocean range switching events possibly enabled by powered flight (Upchurch, et al., 2015). However, such a lack of statistical support for vicariance among pterosaurs might also be attributable to fossil sampling biases. Therefore, a separate, more focused biogeographic analysis of Mesozoic avians, combined with investigation of the dispersal abilities of various modern bird clades, should be the next step in tackling this issue. Despite these uncertainties, our results demonstrate that continental dispersal, extinction, vicariance, and founder-events were important biogeographic processes during coelurosaurian evolution. Major dispersal corridors included the Apulian Route and the Bering Land Bridge, and Europe might have been an important dispersal centre and gateway for coelurosaurs before the mid-Early Cretaceous.

## ACKNOWLEDGEMENTS

We thank Mr. Kenneth HC Fung and the First Initiative Foundation for their support of the International Pennaraptoran Dinosaur Symposium which was the catalyst for this volume chapter. A.D.’s participation in this project was supported by undergraduate student funds from a General Research Fund grant (17103315) awarded to M.P. by the HK Research Grants Council. A.D. was also supported by funding from the HKU MOOC course *Dinosaur Ecosystems* (awarded to M.P.) and from Dr. Jason R. Ali of HKU’s Department of Earth Sciences. Dr. Ali is also thanked for his input in project discussions.

